# Deep Learning Reveals Persistent Individual Signatures in Bat Echolocation Calls of the Greater Leaf-nosed Bat

**DOI:** 10.64898/2026.03.31.715443

**Authors:** Aoqiang Li, Wantao Huang, Xinyuan Xie, Weiming Wen, Liang Ji, Huijun Zhang, Chengyun Zhang, Jinhong Luo

**Affiliations:** Key Laboratory of Pesticide & Chemical Biology of Ministry of Education, Hubei Key Laboratory of Genetic Regulation and Integrative Biology, School of Life Sciences, Central China Normal University, Wuhan 430079, China; School of Electronics and Communication Engineering, Guangzhou University, Guangzhou, 510006, China; Hubei International Science and Technology Cooperation Base for Ecology and Environment, Central China Normal University, Wuhan, 430079, China

**Keywords:** Acoustic communication, Artificial intelligence, Biomarker, Bioacoustics, Wildlife

## Abstract

Intraspecific variation is a prerequisite for natural selection and can manifest in various phenotypic traits, including vocal signals. However, classifying individuals based on their vocalizations, or acoustic individual identification (AIID), remains a significant challenge. This is particularly true for species that use rapidly varying echolocation calls for orientation. Here, we demonstrate that deep learning can overcome the limitation of traditional methods and reveal persistent individual signatures within bat echolocation calls. We recorded echolocation calls from 34 individuals of the greater leaf-nosed bat (*Hipposideros armiger*) under controlled laboratory conditions, with 19 individuals recorded repeatedly over three months. We show that a convolutional neural network (CNN) dramatically outperforms a traditional method, achieving an average identification accuracy of 84% for single calls and 91% for call sequences. In contrast, the traditional Discriminant Functional Analysis method achieved accuracies of only 39% and 47%, respectively. Through systematically altering the temporal structure of echolocation calls in input sequences, we found that temporal patterning enhances individual classification accuracy, suggesting it contributes to the encoding of individual-specific information. This study revealed that echolocation calls of *H. armiger* can contain stable, individual identity that were previously undetectable. Our findings highlight the potential of deep learning for non-invasive AIID and provide a methodological basis for future studies aiming to monitor animals in more dynamic environments.

## 1 INTRODUCTION

Intraspecific variation in genetic traits can provide individuals with adaptive advantages to drive natural selection ^1^. Individual identification serves as the cornerstone of ecological and evolutionary research, providing critical insights into population dynamics, social structures, and movement ecology ^2–4^. To date, individual identification and tracking in the wild have relied heavily on conventional techniques like mark-recapture and telemetry ^5–7^. However, these methods suffer from significant limitations: they are labor-intensive, invasive, and susceptible to observer bias or behavioral alterations in the study subjects ^8–10^.

Bioacoustic signals represent a promising alternative approach for individual identification, given that vocalizations encode individual-specific features through genetic, environmental, and cultural influences ^11–14^. Acoustic individual identification (AIID), which involves identifying individuals by their calls, eliminates the need for making physical contact with the individual, allows efficient long-term monitoring, and reduces personnel costs. Early AIID implementations using traditional machine learning, including support vector machines, Hidden Markov Models, or Discriminant Function Analysis (DFA) ^15–17^, demonstrated conceptual promise but suffered critical constraints: reliance on expert knowledge and experience, vulnerability to ambient noise contamination and inter-class spectral overlap, constrained generalizability under small-sample conditions, and performance degradation caused by low signal-to-noise ratios (SNR) ^18,19^. These limitations critically impair identification fidelity, often resulting in classification accuracy that falls below 50%, which is insufficient for robust ecological inference and restricts researchers to studying individuals only in specific contexts.

Recent advances in deep learning, particularly Convolutional Neural Networks (CNNs), offers paradigm-shifting capabilities for pattern recognition, garnering increased attention from ecologists ^20–22^. Unlike traditional machine learning approaches that rely on hand-crafted feature extraction, CNNs autonomously extract discriminative features through hierarchical learning of spectrotemporal patterns, bypassing error-prone manual feature selection while maintaining robustness to acoustic variability ^23^. Such tools can automate the analysis of various types of data, ranging from species abundance to behavioral patterns and derived from multiple data sources such as images or audio recordings, which are widely used by ecologists for automated species identification, environmental monitoring, ecological modelling or behavioral studies ^24,25^. Moreover, the use of deep learning methods for individual identification has also become a subject of heightened research involving diverse taxa, such as humans, primates, and birds ^26–29^. However, most existing applications focus on image-based individual identification, while the use of acoustic signals for individual identification remains underdeveloped. A key unresolved question is whether deep learning is suitable for AIID, considering the pronounced intra-individual variability in vocal signals in many taxa ^20^.

Bat echolocation calls are well-known for their remarkable plasticity, which vary across behavioral tasks, environments, and time ^30–33^, thus representing a highly challenging case for robust AIID. The high flexibility of their calls has led to enduring scientific controversy over whether bat echolocation calls truly encode individual signatures ^34,35^. Additionally, the physical and ecological demands of echolocation may impose evolutionary constraints on vocal inter-individual variability. Because echolocation calls must maintain acoustic features optimized for target detection and spatial navigation, the extent of inter-individual variability may be more limited than in many non-echolocating species, where variability in vocal signatures is important for individual recognition. Determining whether stable individual signatures can still be extracted under such constraints represents a key challenge for acoustic individual identification in echolocating mammals. Notably, stability does not imply invariance of acoustic signals, but rather the persistence of individual-specific information despite substantial within-individual variability across time and context. Addressing this question would help to reconcile the debate whether individual signatures in echolocation calls exist, and if so, it can facilitate an application of AIID as a powerful tool for the behavior and ecology in a wide range of vocally active species.

Here, we first conducted a systematic literature search of peer-reviewed studies applying deep learning to the acoustic identification of species and individuals, establishing baselines to evaluate the performance of our models. We demonstrate that, comparable to species-level classification, deep learning is a powerful method for AIID under controlled laboratory conditions, achieving up to 91% accuracy even when applied to the challenging domain of bat echolocation calls.

## 2 MATERIALS AND METHODS

### 2.1 Literature survey

We first investigated whether deep learning has been successfully applied to acoustic identification of species and individuals across taxa, and if so, with what accuracy. We conducted a systematic literature search of peer-reviewed studies that applied deep learning to species and individual identification based on acoustic signals. The literature search was performed using the Web of Science "all databases" on February 28, 2025, with the following search terms: TS = “individual identification” AND TS = “deep learn” or TS = “species identification” AND TS = “deep learn”. After screening titles, abstracts, and keywords to exclude irrelevant publications, with particular attention to removing studies based on image-based identification, we retained 107 studies focusing on acoustic identification across multiple taxa, including birds, bats, amphibians, marine mammals, and invertebrates (**Table S1**). For each relevant study, we extracted reported classification accuracy. When data were presented in figures, we used the function “grabit.m” in MATLAB (R2021b, MathWorks, USA) to extract the values ^36^. Furthermore, to compare the performances of AIID between deep learning and non-deep learning methods, we also used the Web of Science “all database” search and performed a search with the following terms: TS = “individual identification” AND TS = “bats” to find studies on bat acoustic individual identification based on traditional methods, which yielded 14 studies (**Table S2**). We extracted the classification accuracy (*A*) and sample size (*N*), and additionally calculated the chance level or baseline chance accuracy (*C*) and the normalized accuracy (Cohen’s Kappa, *κ*) following equations:

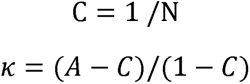

### 2.2 Echolocation call recording

Thirty-four adult the greater leaf-nosed bats (*Hipposideros armiger*) were used for acoustic individual identification (**Figure 2A**). These bats were captured from two geographical locations separated by approximately 900 km. Fifteen individuals were caught from a cave located in Hanzhong County, Shaanxi Province, China and transferred to the housing facility in Northeast Normal University (NENU). Nineteen individuals were caught from a cave located in Xianning County, Hubei Province, China and transferred to the housing facility in Central China Normal University (CCNU). Bats were kept in temperature and humidity regulated rooms with 12-hour light-dark cycles. Bats had *ad libitum* access to water and food. Capture, housing and experimental protocols were approved by the Institutional Animal Care and Use Committee of the respective university (Permits: NENU-2018-03 and CCNU-IACUC201902191427).

We recorded the echolocation calls of individual bats hanging from a perch in echo-attenuated rooms, with the walls and ceiling covered by thick acoustic foams and the floor by a nylon blanket ^37–39^. At CCNU, recordings were conducted in a flight room (5 × 4 × 2.2 m) (**Figure S1A**). Individual bats were recorded while hanging at a tripod approximately 1.5 m above the floor. A custom-made ultrasonic microphone (based on SPU0410LR5H, Knowles Corporation, Itasca, IL, USA), approximately 50 cm from the bat, was directed toward the bat. The microphone signals were amplified via microphone preamplifiers (OctaMic II, RME, Haimhausen, Germany) and sampled with au audio interface (Fireface 802, RME, Germany) at a rate of 192 kHz. The bats were recorded multiple times per week over three months. At NENU, the recordings were provided by our collaborators and were originally collected for a different experimental purpose. Echolocation calls were recorded at a sampling rate of 250 kHz with an array of 16 ultrasonic microphones (NEUmic, Ultra Sound Advice, UK) using a multichannel data acquisition board (PXIe 6358, National Instruments, USA) via customer LabVIEW programs (**Figure S1B**). The recordings were conducted in a flight room of 7 × 4.8 × 2.6 m size and the bat hang from a perch attached to the ceiling and positioned 30 cm below it. The microphones were placed on the floor and pointed towards the bat. The bats were recorded for five days in total. The recording time lasts for 5 seconds. A comprehensive summary of the dataset is shown in **Table S3**. To combine the two recording datasets, recordings sampled at 250 kHz were down-sampled to 192 kHz before further processing.

### 2.3 Call sequence segmentation

This study utilized two distinct datasets derived from labeled audio recordings. The first was composed of standard 5-second call sequences, referred to as sequence-level identification. The second dataset consisted of single calls, which were uniformly zero-padded to 1 second to maintain dimensional consistency, referred to as call-level identification. The single calls were obtained by processing call sequences through an Endpoint Detection Algorithm. Briefly, the call sequences firstly underwent several preprocessing steps, including the removal of the DC component and amplitude normalization, which collectively serve to enhance the stability and robustness of subsequent signal analysis and feature extraction. A Hamming window (length = 200 points) was employed for audio framing, with adjacent frames overlapping by 120 points (60% overlap ratio) to maintain signal continuity. Following the framing process, energy-entropy ratio features were extracted, with high and low thresholds set to 0.1 times and 0.05 times the dynamic range, respectively. To mitigate transient noise, a 10-point median filtering technique was employed. Furthermore, dynamic thresholds were established by analyzing a leading 0.25-second silent segment, combined with constraints of a minimum valid segment (5 frames) and maximum silent interval (8 frames), to precisely extract individual vocalization clips from each sequence.

### 2.4 Discriminant function analysis

Discriminant function analysis (DFA), a statistical classification method designed to identify optimal linear combinations for distinguishing between two or more predefined groups, was implemented for individual identification. We selected DFA due to its historical prevalence in bat acoustic studies, allowing for a direct reference to prior literature. Two distinct models for bat individual identification were constructed, utilizing call sequence datasets and single call datasets, respectively. For call-level identification, the frequency spectrum surrounding the dominant second harmonic of the echolocation calls (60–80 kHz) was firstly transformed to the Mel scale via a Discrete Fourier Transform (DFT), which provide a nonlinear resampling of the frequency axis and a compact representation of spectral information for machine-learning analysis. Mel-frequency representations were chosen due to their widespread application in the machine learning community. Although the Mel scale is essentially a logarithmic transformation that de-emphasizes higher frequencies, restricting the frequency range to 60–80 kHz effectively preserves the spectral detail relevant to these echolocation calls. Then, feature extraction employed a Discrete Cosine Transform (DCT) to derive Mel-Frequency Cepstral Coefficients (MFCCs), Zero Crossing Rate (ZCR), and Spectral Centroid (SC). Mean and standard deviation values of these features were calculated to generate 44-dimensional feature vectors (comprising 40 dimensions from 20 MFCCs, 2 dimensions from ZCR, and 2 dimensions from SC), ensuring stability for subsequent model training and classification. For individual identification, Linear Discriminant Analysis (LDA) was employed using Python’s librosa library with the following parameters: no pre-emphasis, n_mfcc=20, frame size=2048 samples, hop length=512 samples, and Hann window spectral smoothing. This pipeline maintained dimensional consistency across training and test sets while capturing stable acoustic characteristics for classification. During analysis, the total dataset was partitioned with 10% allocated as the test dataset, while the remaining 90% was divided into training and validation datasets through 5-fold cross-validation. The same methodology was applied to the sequence-level identification.

### 2.5 Deep learning

The experiment uses the same datasets as the DFA model for comparative analysis, with experimental conditions grouped into two main experiments: call- and sequence-level identification. To investigate whether deep learning-based individual identification models rely on spectral features and temporal cues. At the call-level, the following conditions were included: single call (original single call datasets), multiple consecutive calls (sequences of 2 to 23 calls were extracted in their natural, original order from the call sequences dataset), and multiple randomly selected calls (2 to 23 calls randomly sampled from different positions within the call sequences dataset, with no regard to their original order). For the sequence-level, the conditions included: 5-second call sequences (original single call datasets), 5-second time-reversal sequences (reversing the order of calls within the sequences), 5-second position-random sequences (calls within the sequences randomly reordered, disrupting natural temporal structure), and call-swapping sequences (hybrid sequences constructed by replacing the spectral content of one individual’s calls with that of another, while preserving the original temporal structure). By comparing classification performance across these conditions, we tested three hypotheses. First, if temporal cues contribute to encode individual identity, disrupting their natural order of calls should reduce classification accuracy. Second, if spectral features are the primary carriers of identity information, preserving an individual’s spectral content while altering its temporal structure should still yield high accuracy. Conversely, if temporal patterns dominate, sequences with preserved temporal structure but foreign spectral content should show better performance. Third, the call-swapping experiment was designed to directly evaluate the relative contribution of spectral and temporal features to individual identification. For all conditions except single calls and 5-second call sequences, we employed endpoint detection using a ratio method to identify the calls in each audio file, obtaining the starting and ending frames for each call. We performed echo filtering and saved each call individually, when the interval between the previous call and the current call is less than 0.01 seconds and the maximum amplitude of the previous call is more than three times that of the current call.

For individual identification, each condition firstly was partitioned with 10% allocated as the test dataset, while the remaining 90% was divided into training and validation dataset through 5-fold cross-validation. Each audio segment underwent Short-Time Fourier Transform (STFT) with a 4096-point Fourier window and 2048-point frame shift to compute time-frequency spectrograms. A bandpass filter (55 to 85 kHz) was applied to retain frequency bands containing the dominant second harmonic component. Concurrently, the spectrograms were converted to log-Mel spectrograms using 128 Mel filter banks for the 60-80 kHz frequency range to achieve dimensionality reduction ^40^. The ResNest50d network was selected as the backbone for feature extraction and classification ^41^. This choice was motivated by three main considerations ^42,43^. First, CNN architectures with integrated attention mechanisms have been shown to substantially improve classification performance in bioacoustic contexts, where features are subtle and background noise is complex. However, it is worth noting that the split-attention mechanism in ResNeSt operates on grouped features along the channel dimension rather than on spatial positions within the input spectrogram. As a result, the attention weights do not directly indicate which time–frequency regions are most influential, limiting the direct interpretability of the model’s decisions in bioacoustic terms. Second, empirical evidence indicates that ResNeSt50 outperforms other mainstream CNN architectures when processing complex animal vocalization spectrograms, demonstrating its strong ability to capture intricate time–frequency patterns. Third, the ResNeSt50d variant incorporates architectural enhancements from the ResNet-D design, enabling the model to reduce feature map dimensionality while preserving the structural integrity of acoustic signals. This helps minimize the loss of key discriminative information, which is crucial for individual-level identification in bat echolocation calls. Nevertheless, we also selected EfficientNet-B0 as the backbone network for feature extraction and classification to provide a more comprehensive evaluation. During the training process, the model parameters are iteratively updated using a Stochastic Gradient Descent (SGD) optimizer with momentum, with an initial learning rate set to 0.01, a minimum learning rate of 0.0001. A cosine annealing learning rate scheduling strategy was implemented, where the annealing cycle *T_max_* was set to 100, matching the total training epochs. To explore the acoustic basis of individual signatures, we separated each echolocation call into its constant frequency (CF) and frequency-modulated (FM) components and performed individual recognition on each component independently. The CF and FM components were separated by estimating the end of the CF component and the start of the FM component, using custom-written scripts in MATLAB ^44,45^. We trained our models on an NVIDIA RTX 4090 GPU over 100 epochs with batch size of 32 and binary cross-entropy based on focal loss.

### 2.6 Statistical analysis

All statistical analyses were conducted using R Studio (v 2022.10.0+353, R version 4.2.2) and MATLAB (R2021b, MathWorks, USA). Principal component analysis was used to examine the variability of individual calls using the *FactoMineR* package based on acoustic parameters, including peak frequency, call duration, bandwidth, and pulse interval. We employed the Bhattacharyya coefficient to quantify the degree of overlap between individuals’ call distributions in the reduced feature space. A higher coefficient value indicates greater overlap, meaning that individuals are difficult to distinguish based solely on these linearly transformed feature. We also employed Mann-Whitney U test (as data normality was rejected according to Shapiro-Wilk test in the package *stats*) to compare the differences in average accuracy between groups.

## 3 RESULTS

### 3.1 Accuracy for acoustic identification of animal species by deep learning

We conducted a comprehensive review of 110 studies that evaluated acoustic identification of species based on deep learning. These studies encompassed a broad range of species, from well-studied taxa such as birds (67.29%) and bats (20.56%) to less frequently studied amphibian, marine mammals, and invertebrates (**Figure 1A; Table S1**). Regarding the classification performance, the overall average identification accuracy for all taxa was 0.89 ± 0.08 (mean ± SD), with the highest average accuracy for marine mammals at 0.92 ± 0.03 (mean ± SD). For bats, the average species-level identification accuracy was also high at 0.89 ± 0.08, and the lowest for invertebrates at 0.86 and was derived from only a single study (**Figure 1A**). Additionally, we systematically reviewed 17 studies focusing on the bats acoustic individual identification based on traditional methods. We found the identification accuracy for individual bats based on traditional methods was much lower, at 0.48 ± 0.22 (mean ± SD), with the normalized accuracy of 0.46 ± 0.23 (mean ± SD; **Figure 1B; Table S2**).

**Figure 1.**
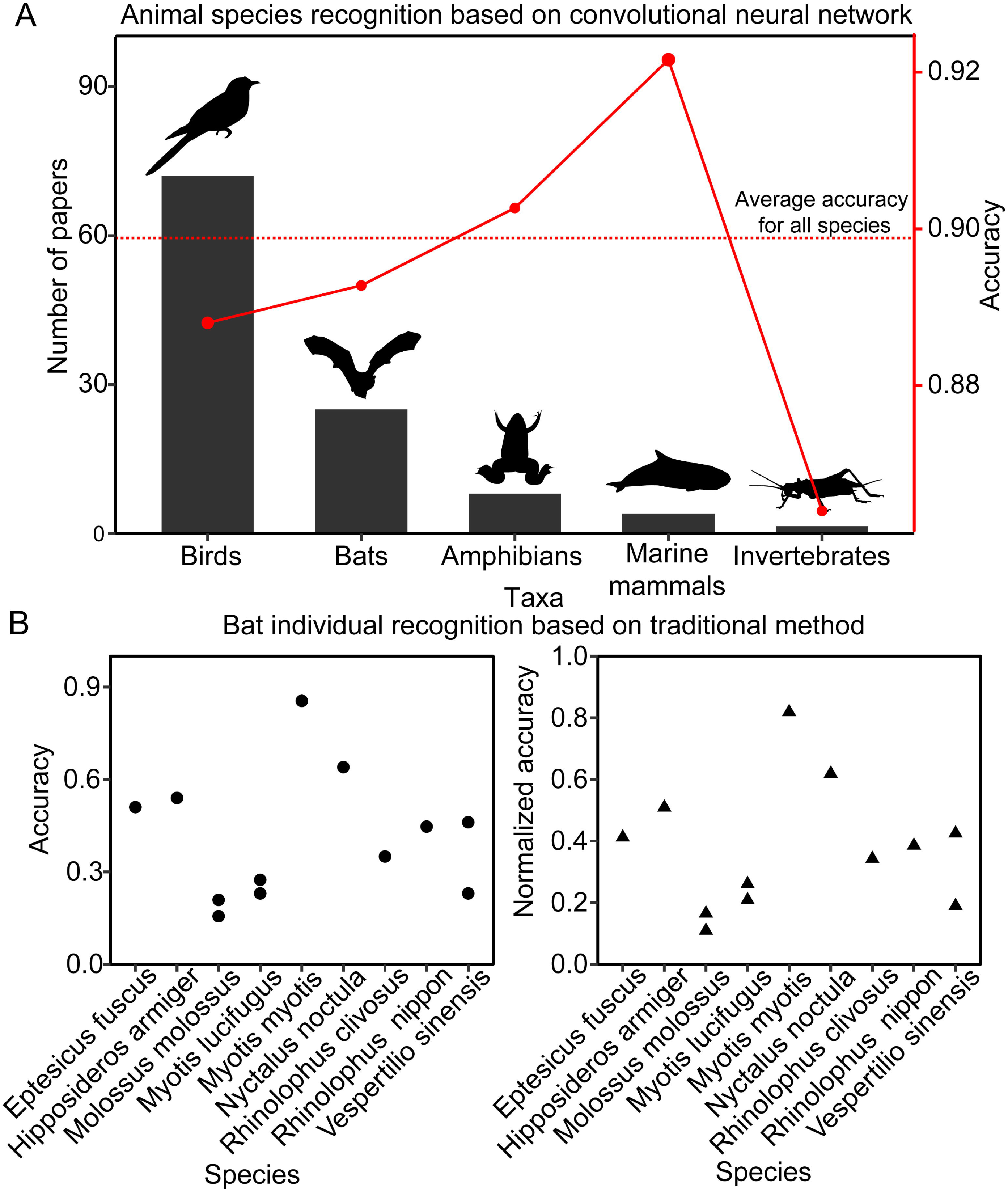
Acoustic species identification in representative taxa and acoustic individual identification in echolocating bats. (A) Acoustic species identification across taxa using convolutional neural networks revealed an average accuracy of 0.89 ± 0.08 (mean ± SD). (B) All previous researches on acoustic individual identification in bats have used traditional machine learning methods, with an average accuracy of 0.48 ± 0.22 (mean ± SD).

### 3.2 Substantial intra-individual variability in bat echolocation call across time

Apart for the individuals from NENU, we conducted three-month acoustic recording of *H. armiger*, yielding a total of 17,584 standardized 5-second vocalization sequences. **Figure 2B** displays an example of the waveform of a 5-second call, as well as the waveform and spectrogram of a single call. We found that call parameters such as peak frequency, duration, bandwidth, and pulse interval varied considerably across different recording days, indicating that even for the same individual, there is a high level of variability in multiple call features (**Figure 2C**). Principal component analysis further revealed that most calls clustered together, with an average Bhattacharyya coefficient, which is a measure of the amount of overlap between two statistical samples or populations, of 0.80 ± 0.14 (mean ± SD) for 20 individuals recorded on the same day, suggesting limited acoustic distinctiveness among individuals (**Figure 2D**). Similar results were obtained for all individuals.

**Figure 2.**
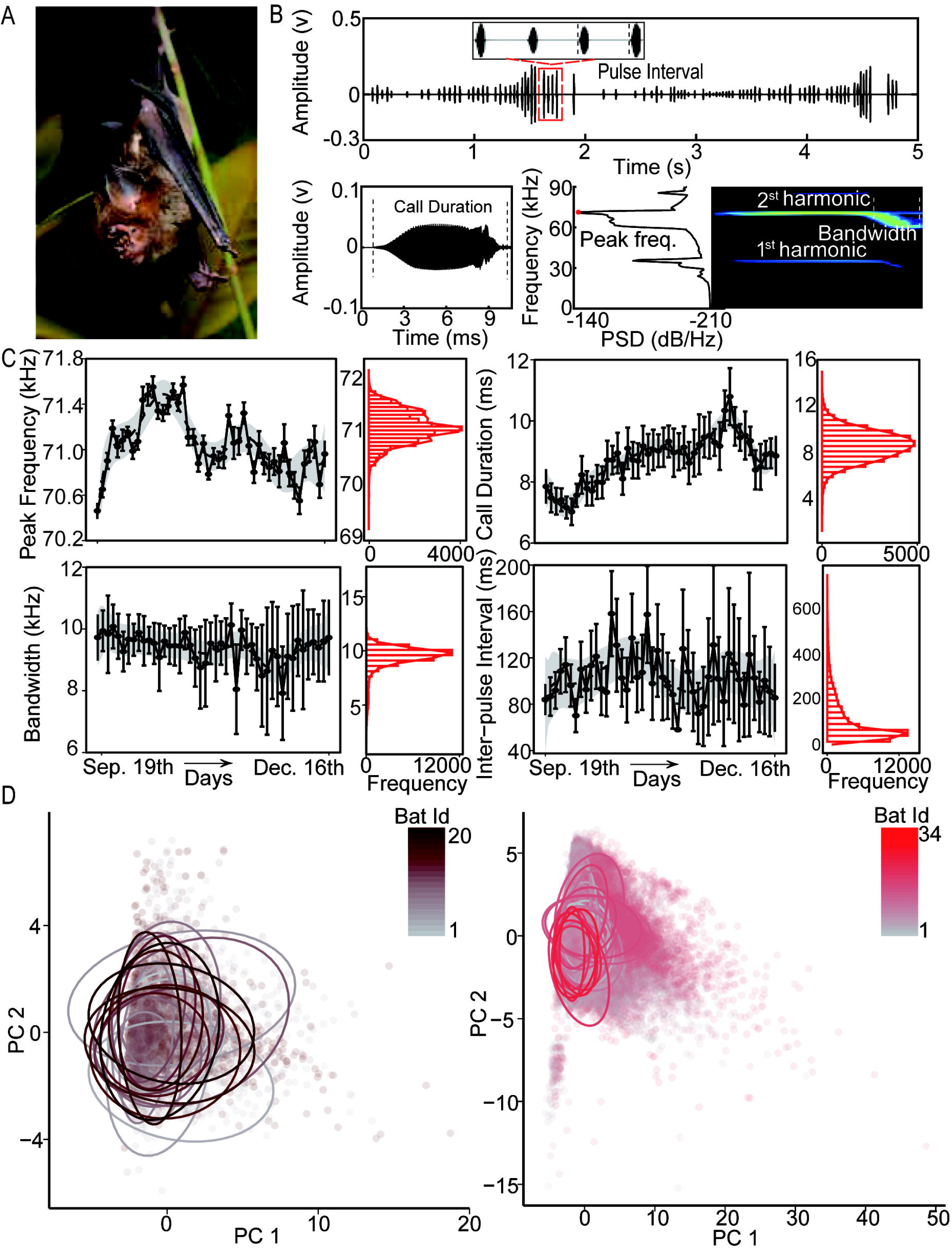
Intra-individual vocal variability in Hipposideros armiger under controlled laboratory conditions. (A) A portrait of H. armiger. (B) Examples of the echolocation call. The top panel displays the waveform of a 5-second echolocation sequence, while the bottom panel shows the waveform of a single call alongside the corresponding spectrogram and power spectrum. (C) Illustrations of daily variation in four call parameters (peak frequency, duration, bandwidth, and pulse interval) measured from one individual, representative of the general pattern observed across all individuals. (D) Principal Component Analysis (PCA) of 20 individuals recorded on the same day (left panel) and 34 individuals recorded across multiple days (right panel), with ellipses representing 95% confidence intervals. The degree of overlap between individuals’ calls was quantified by the Bhattacharyya coefficient, where a higher value indicates greater overlap.

### 3.3 AIID by the traditional DFA

We first employed traditional methods to investigate AIID of bats. The results of the DFA showed that the average classification accuracy based on single call for the 34 individuals was 0.39 ± 0.21. Furthermore, no significant difference in classification accuracy was observed between the CCNU and NENU sites (Mann-Whitney U test: *P* = 0.19; **Figure 3A**). For 5-second call sequences recognition, the average classification accuracy improved to 0.47 ± 0.26, with classification accuracy at the CCNU site significantly higher than that at NENU (Mann-Whitney U test: *P* = 0.02).

**Figure 3.**
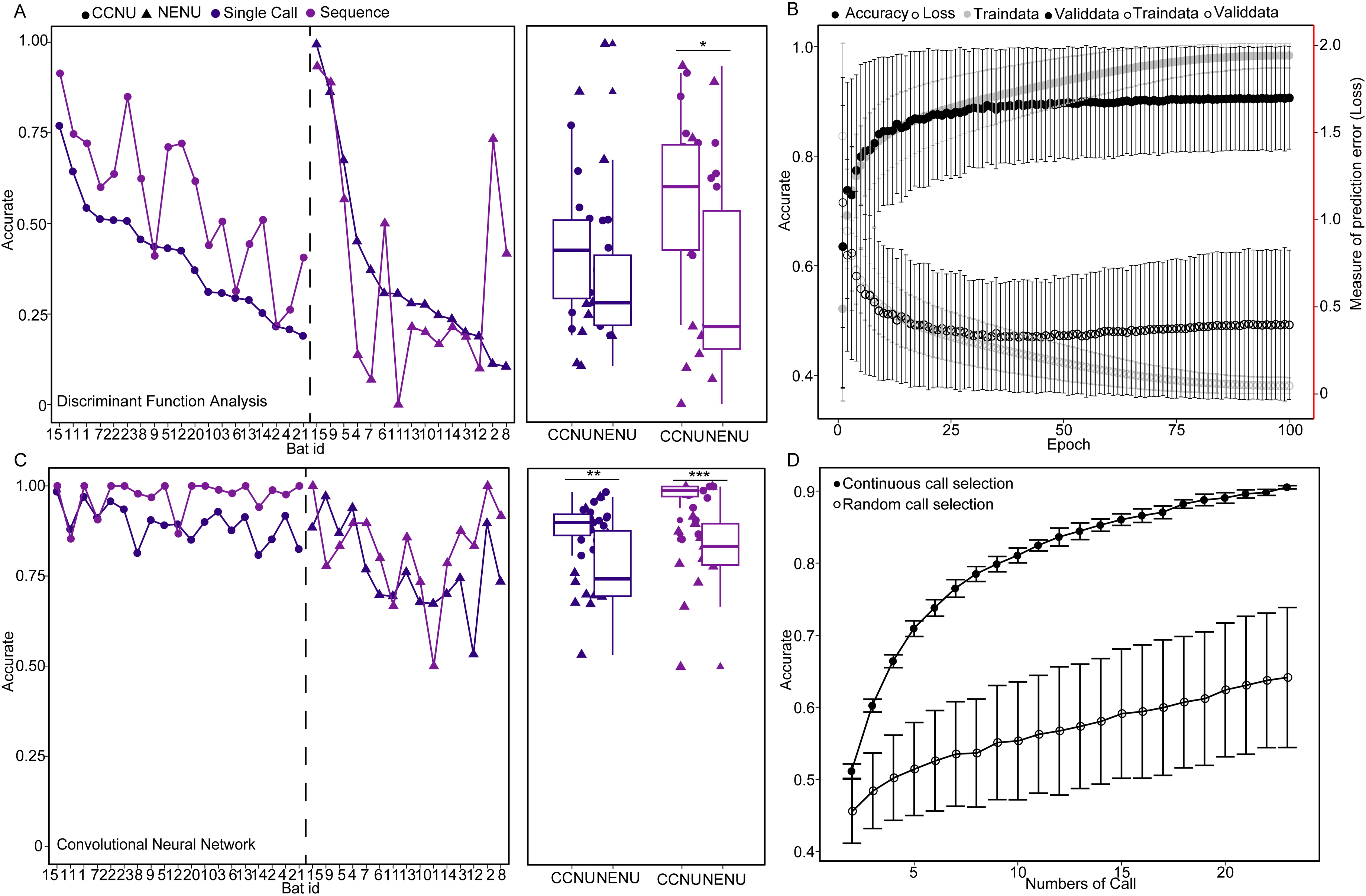
Comparison of acoustic individual identification performance in bats using traditional methods versus deep learning. (A) A comparison of classification accuracy between CCNU and NENU based on DFA for single-call and 5-second call sequence paradigms; (B) Changes in accuracy and loss for both the training dataset and validation dataset as epochs progress (mean ± sd); (C) A comparison of classification accuracy between CCNU and NENU based on deep learning for single-call and 5-second call sequence paradigms; (D) The variation in accuracy as the number of selected calls changes, with solid circles indicating consecutively selected calls and hollow circles representing randomly selected calls (mean ± sd).

### 3.4 AIID by deep learning

In contrast, deep learning achieved markedly higher performance. The model showed stable convergence throughout training, with classification accuracy consistently increasing and cross-entropy loss decreasing as epochs progressed (**Figure 3B**). Although classification accuracy varied among individuals based on single call, the majority exhibited relative high classification accuracy (> 0.50), achieving an average classification accuracy of 0.84 ± 0.11 (mean ± SD) and the normalized accuracy of 0.84 (**Figure 3C and Figure S2A**). Notably, accuracy at CCNU was significantly higher than at NENU (0.90 vs. 0.77: Mann-Whitney U test: *P* = 0.002). Furthermore, we observed a progressive increase in classification accuracy with the number of calls incorporated into the analysis. Multiple consecutive calls consistently outperformed those constructed from multiple randomly selected calls (**Figure 3D**), suggesting temporal sequence information may help to enhance classification performance.

Compared to single-call analysis, the classification accuracy improved for most individuals under the 5-second call sequence paradigm, culminating in an overall mean accuracy of 0.91 ± 0.12 (mean ± SD) and the normalized accuracy of 0.91 (**Figure 3C**). The classification accuracy at CCNU was also significantly higher than at NENU (Mann-Whitney U test: *P* < 0.001), reaching 0.97 compared to 0.83. When using EfficientNet-B0 as the backbone network, the mean accuracy showed a slight decrease but remained at 0.89 ± 0.10, and the classification accuracy at CCNU remained higher than that at NENU (0.94 vs. 0.84: Mann-Whitney U test: *P* = 0.02; **Figure S2B**).

**Figure 4A** displays the schematic representation of 5-second time-reversal sequences and 5-second position-random sequences. We observed that both time-reverse and position-random calls led to statistically significant reductions in accuracy compared to natural sequences (Mann-Whitney U test, all P < 0.01; **Figure 4B**), while no significant difference was observed between these two modifications (Mann-Whitney U test; P = 0.14), suggesting that temporal cues play a key role in encoding individual identity and that maintaining natural call order is critical for accurate recognition. Besides, **Figure 4C** displays the schematic representation of call-swapping sequences. We found that when synthesizing novel call profiles through recombination of spectral features and temporal sequence information, individuals identified based on spectral features (feature dominated) exhibited significantly higher accuracy than those identified from temporal sequence information (position dominated) (**Figure 4D**). This indicates that the importance of spectral features in recognition is greater than that of temporal sequence information. However, when each call was decomposed into its CF and FM components and evaluated independently, the classification accuracy deteriorated markedly (CF-only: 0.34 ± 0.28; FM-only: 0.30 ± 0.39; **Figure S3**).

**Figure 4.**
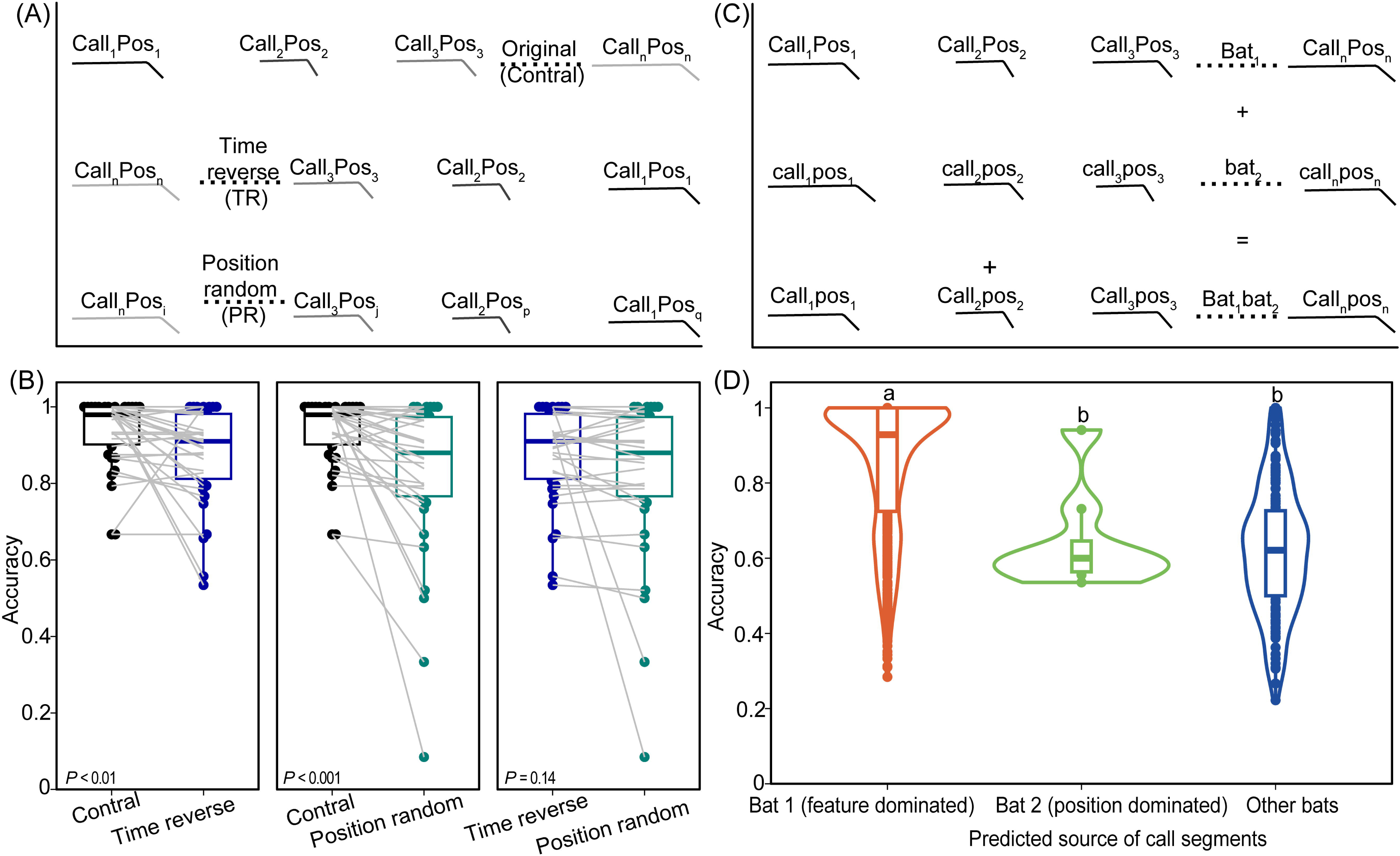
Exploring the importance of spectral features and temporal sequence information in individual identification. (A) Schematic representation of 5-second time-reversed (reversing the order of calls within the sequences) and randomly repositioned call sequences (calls within the sequences randomly reordered, disrupting natural temporal structure), illustrating the experimental design; (B) Comparison of accuracy among different groups; (C) Schematic representation of call-swapping sequences (hybrid sequences constructed by replacing the spectral content of one individual’s calls with that of another, while preserving the original temporal structure), demonstrating the approach used for synthesizing novel call profiles through the recombination of spectral features and temporal sequence information. (D) Comparison of accuracy among different groups. The numbers represent the corresponding identification counts.

## 4 DISCUSSION

In this study, we successfully applied deep learning to AIID under controlled laboratory conditions and achieved an unprecedented recognition accuracy of up to 0.91, which is comparable to the benchmark for species identification (0.89 on average), despite the inherent challenge of discerning minute individual vocal differences. This achievement is particularly striking given that individual identification demands resolving acoustic variations orders of magnitude smaller than between-species distinctions. In contrast, traditional methods, such as discriminant function analysis (DFA), achieved accuracy approximately two times lower than that of the deep learning, underscoring deep learning’s capacity to decode nonlinear spectrotemporal interactions impervious to manual feature engineering. Our dataset, comprising 34 bats monitored over three months and 17,584 calls, demonstrates the natural range of vocal plasticity in bat echolocation calls. This variability, driven by factors such as environmental noise ^46^, behavioral states ^47^, and ontogenetic changes ^48^, and social factors ^49^, has historically posed challenges for conventional methods ^34,50^.

The primary objective of this study was to establish a proof-of-concept under controlled conditions, specifically to evaluate whether deep learning is suitable for AIID. Our results suggest that deep neural networks effectively leverage hierarchical feature abstraction to isolate stable ‘vocal fingerprints’, even in groups with excessive intra-individual variability in acoustic signals, achieving robustness where traditional methods tend to struggle ^51^. Although individual call features such as frequency or duration vary substantially across time, the relative configuration and joint distribution of these features appear to form a robust identity signature. Deep learning models are particularly well suited to capture such distributed representations, enabling reliable identification even when individual acoustic components are highly variable. In this context, our use of “reveals” refers to the model’s ability to detect and utilize identity-relevant information, rather than to explicitly identify the precise acoustic features involved. However, the noticeably higher classification accuracy at CCNU compared to NENU suggests that site-specific factors may have influenced model performance, potentially arising from acoustic artifacts introduced by differences in recording environments and equipment, coupled with a lower signal-to-noise ratio (SNR) in the NENU recordings.

Our research highlights the crucial role of spectral features in individual identification, providing compelling evidence for the encoding of individual information in the echolocation calls of bats. Spectral features, such as frequency and bandwidth of echolocation calls, serve as distinctive biological markers that allow researchers to differentiate between individuals within a species ^52–54^. Studies have shown that variations in the frequency components of echolocation calls can reflect individual identity, with certain bats exhibiting unique call characteristics that can be consistently recognized across different contexts ^55,56^. Furthermore, the application of advanced bioacoustic analysis techniques ^30,38,57^ has enabled researchers to automate the identification process by analyzing these spectral features, thereby improving the efficiency and accuracy of individual recognition. The encoding of individual labels through spectral characteristics is not merely an analytical tool, but a critical aspect in the broader context of understanding bat ecology and behavior. To further investigate the acoustic basis of individual signatures, we conducted a preliminary analysis by separating each echolocation call into its CF and FM components and performing recognition on each independently. Both CF-only and FM-only models yielded markedly lower accuracy (with averages below 35%), far below the 91% achieved with full-spectrum calls. This suggests that individual identity is not encoded in isolated acoustic components, but rather distributed across the full spectrotemporal structure. This may also explain the poor performance of traditional approaches such as DFA, which rely on manually extracted features. However, we believe this result should not be taken as direct evidence that FM bat echolocation calls cannot encode individual signatures, for several reasons. First, we did not specifically fine-tune the model to optimize performance; therefore, these results likely represent an underestimation of the potential accuracy. Second, the FM component in CF-FM bats appears more stereotyped than in purely FM bats. For instance, both the duration and bandwidth of the FM component in *H. armiger* exhibit a much narrower dynamic range compared to those of many FM bats, such as the big brown bat ^38,58^. Third, there is solid evidence that the echolocation calls of FM bats contain strong individual signatures ^52,54^, although the number of individuals involved in previous analyses has typically been limited. Furthermore, it is critical to interpret these findings within the constraints of the methodology. The deep learning model identifies a robust acoustic signature that could facilitate individual identification. This demonstrates the information encoding capacity of the calls but does not constitute evidence that bats perceptually discriminate or cognitively utilize these cues. The fundamental question of whether bats use these specific, model-derived features for social recognition remains unanswered.

In addition to spectral features, when both spectral and temporal features are integrated, the classification performance improves significantly, highlighting the value of temporal sequence information in individual recognition tasks. This temporal pattern provides context and relational dynamics that are essential for distinguishing between individuals, particularly in species where vocalizations are complex and varied ^59,60^. Thus, the encoding of individual labels through time-based characteristics is not just an auxiliary feature; it is a fundamental element that shapes how organisms interact and communicate within their species ^61^. By incorporating temporal pattern into identification frameworks, such as Recurrent Neural Networks ^62^, researchers can better understand social structures, mating systems, and communication strategies in various species, lending insight into behavioral ecology and evolution. However, it is worth noting that this study is that all recordings were obtained from perching individuals. In many bat species, the timing of echolocation pulses during flight is partially coupled with the wingbeat cycle, which can influence call emission patterns and temporal structure ^63^. Consequently, the importance of pulse interval as a discriminative feature identified in our analysis may differ under flight conditions.

This study establishes bats as model organisms for advancing AIID development, where by surmounting traditional methods limitations through deep learning, our framework enables non-invasive individual studies at under controlled laboratory conditions. These results show great promise for extrapolating to other acoustically active taxa. Echolocation signals are primarily under stabilizing selection for reliable echo detection, which should theoretically minimize individual variation. In contrast, many other animal vocalizations are under diversifying selection to encode information such as individual identity, group membership, or reproductive status. With appropriate adaptation to species-specific acoustic characteristics, such methods may be extended to other vocally complex systems (e.g., cetaceans and songbirds), enabling novel insights into adaptive signal evolution, dynamic social network topologies, host-pathogen coevolution, and anthropogenic impact assessments ^4^.

Nevertheless, application of AIID to freely-ranging wild animals still faces considerable challenges. First, our experiments were conducted on recordings of single bats in a resting state under controlled laboratory conditions. In real ecological environments, bats often fly and move in groups. In such cases, the complexity of acoustic data will significantly increase, potentially leading to a decrease in identification accuracy. Moreover, echolocation calls produced across different behavioral contexts, such as flight, foraging, or prey capture, may exhibit context-dependent acoustic variation driven by task-specific sensorimotor demands, which could further influence the performance of CNN-based individual identification. In addition, environmental complexities such as reflections and reverberation in cluttered habitats and calls from conspecific and heterospecific individuals can introduce spectral-temporal distortions, which require advanced algorithms for separating overlapping calls^64^. A further limitation is that the present study employed a closed-set design, in which all individuals included in the test set were also represented in the training set. In contrast, real-world applications, such as long-term acoustic monitoring of wild animals, will inevitably involve open-set scenarios, where calls from previously unobserved individuals must either be identified as unknown or incorporated into the recognition system. The closed-set framework adopted here therefore represents an initial step toward validating the feasibility of deep learning-based individual identification under controlled conditions. Extending this approach to open-set recognition will be an important direction for future research, potentially involving strategies such as novelty detection or incremental learning to enable robust performance in more realistic ecological settings. Future research should focus on optimizing algorithms, expanding datasets, and enhancing the robustness of models to evaluate whether individual signatures remain detectable across diverse behavioral contexts and ecological environments, a prerequisite for applying AIID to freely behaving wild populations.

## Author contributions

AL, HZ performed the literature survey; AL analyzed the survey data; WH, XX, WW conducted the classification tasks; AL and JL drafted the manuscript; CZ, JL supervised the study.

## Acknowledgements

We thank Manman Lu and Xinyao Yang for collecting the recordings, Aiqing Lin and Jiang Feng for permitting us to use the recordings from NENU, and Jianan Ding and Xuejiao Qin for their valuable discussions. We are grateful to Longgang Pang for performing the initial individual classification tests, which motivated this study. This work was supported by the National Natural Science Foundation of China (32270535 and 32400395) and the China Postdoctoral Science Foundation (30205202371).

## Data availability statement

The datasets and Rcode during the current study are available in the Figshare repository (10.6084/m9.figshare.19613934).

## Conflict of interest statement

The authors declare no competing interests.

---

**Figure S1.**
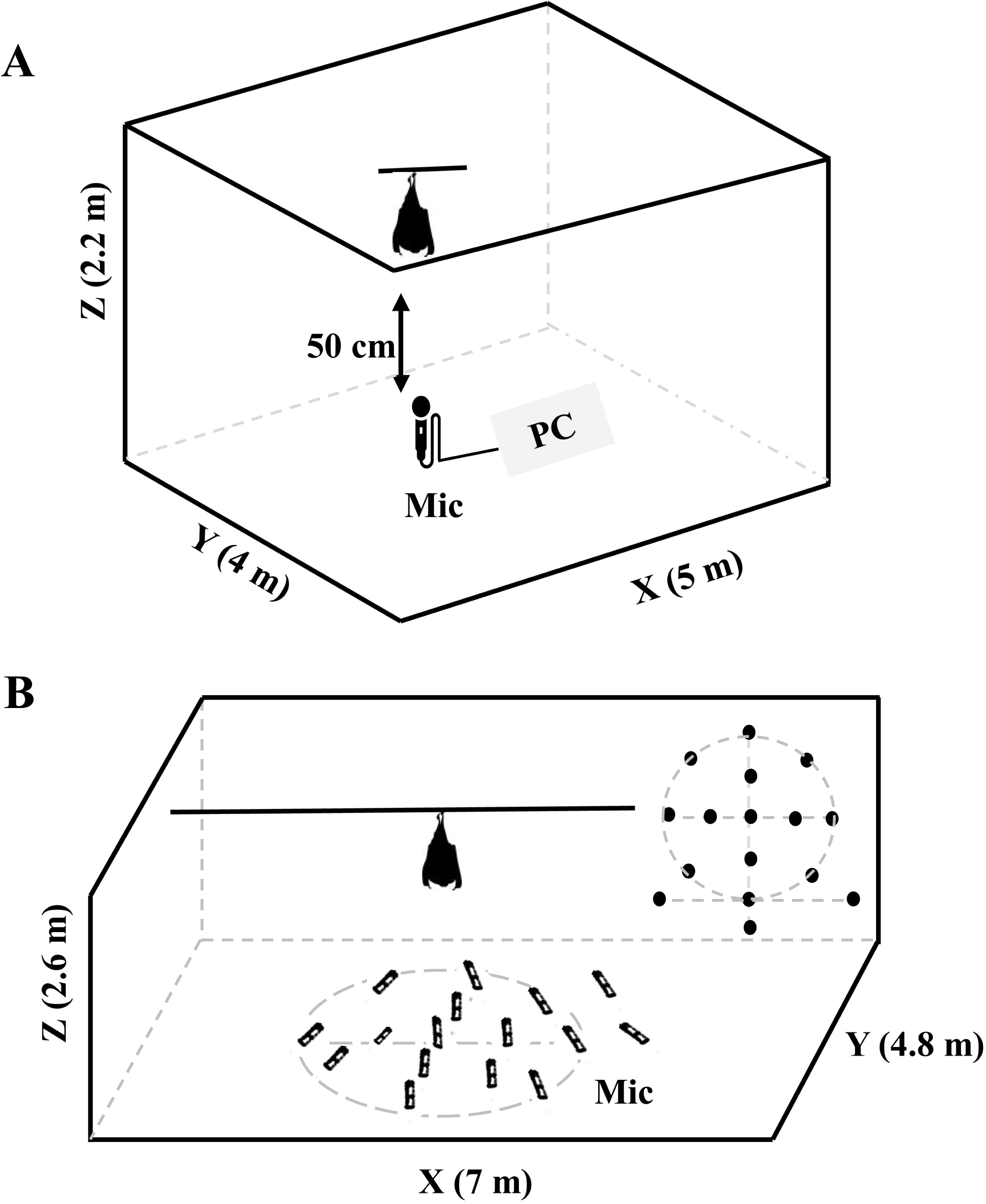
Schematic diagrams of the recording setups at CCNU (A) and NENU (B).

**Figure S2.**
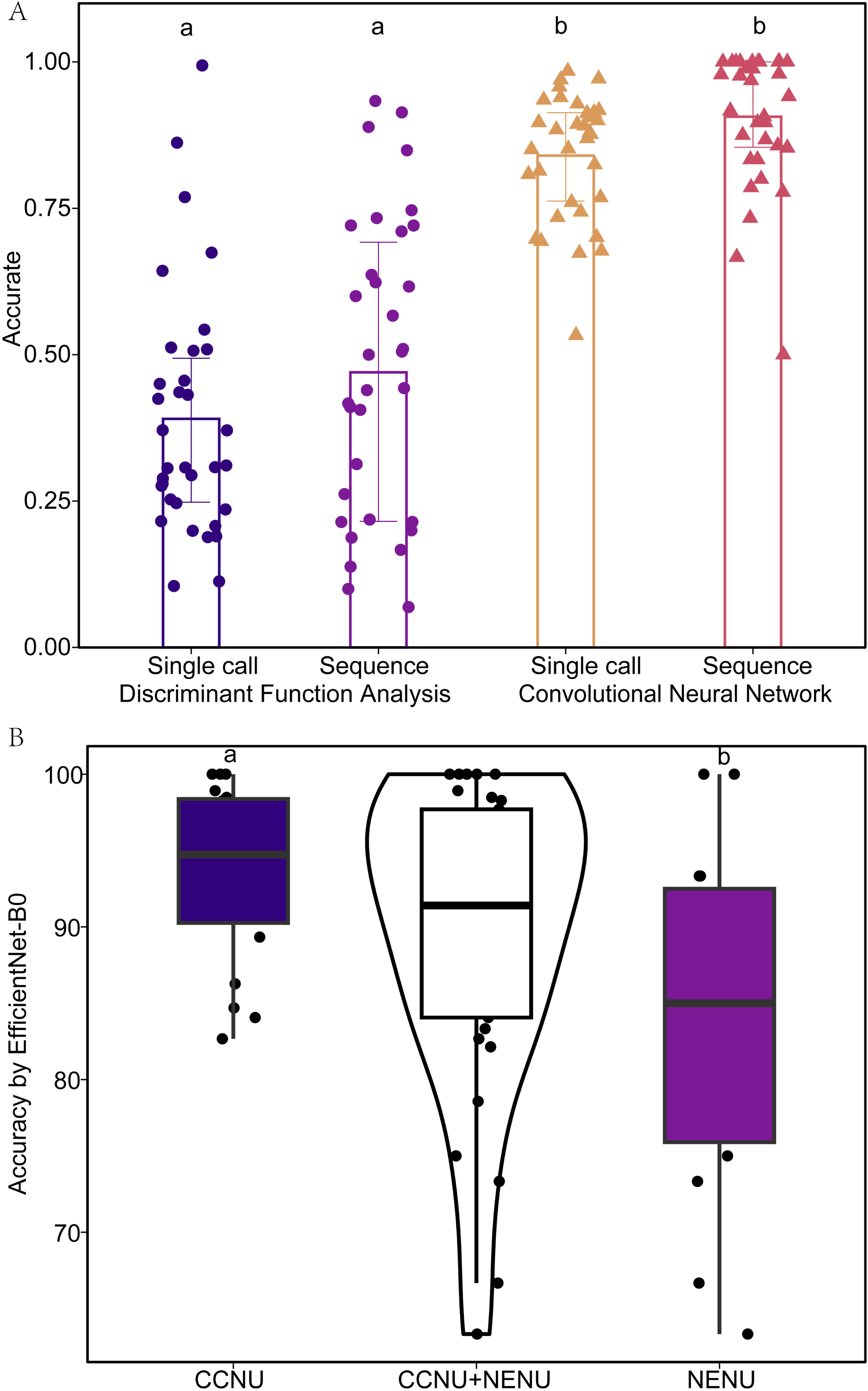
A histogram illustrating the classification accuracy of DFA and deep learning across single-call and 5-second call sequence paradigms based on ResNest50d network (A), and a comparison of classification accuracy between CCNU and NENU based on EfficientNet-B0 network for 5-second call sequence paradigms (B). The center represents the overall classification accuracy, while the two sides indicate the classification accuracy at CCNU and NENU, respectively. Shared letters indicate no significant difference, and distinct letters indicate significant differences based on the Mann-Whitney U test.

**Figure S3.**
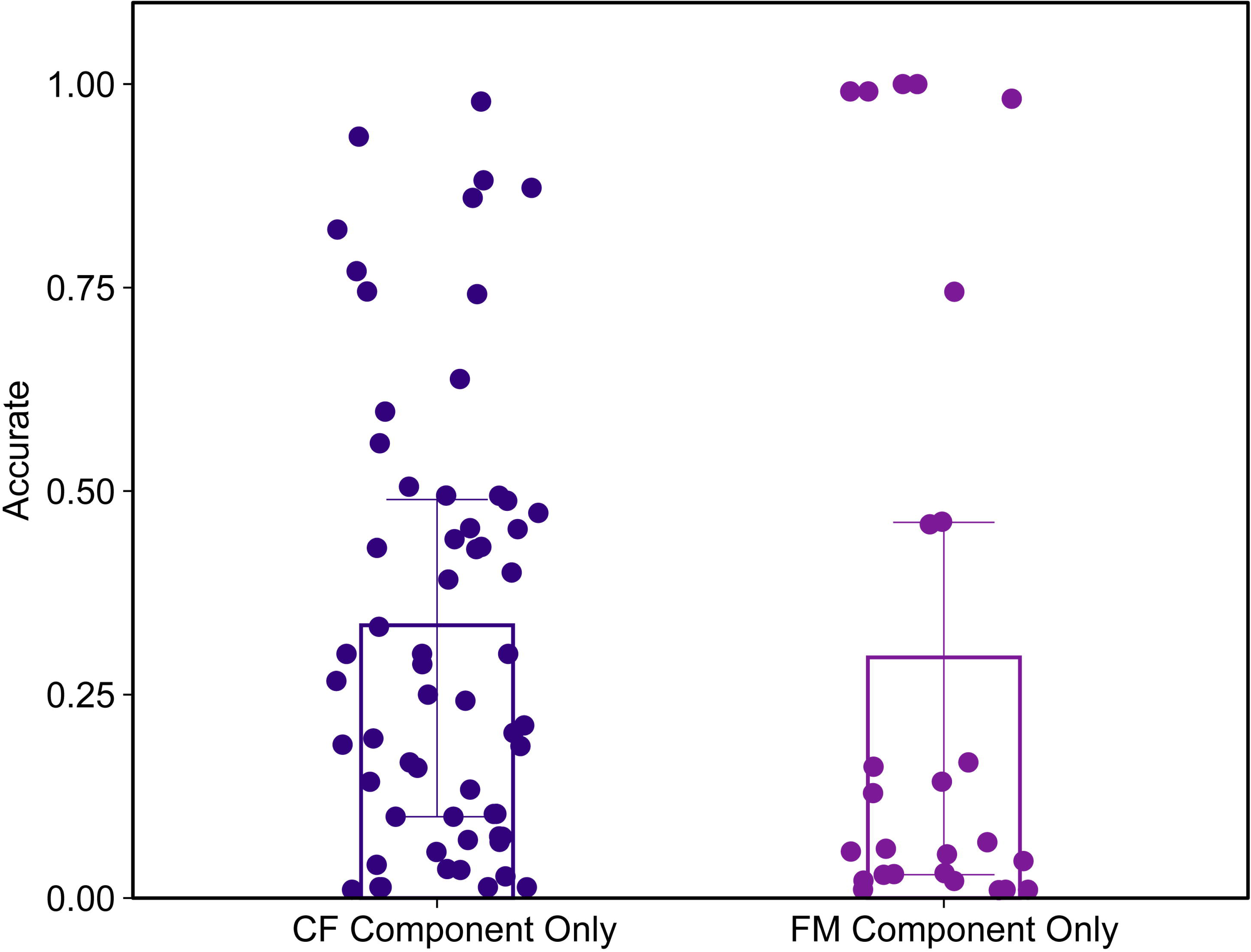
Classification accuracy for CF and FM components alone based on ResNest50d network, with 5-second sequence as inputs.

## Notes

### Competing Interest Statement

The authors have declared no competing interest.

